# The maize heterotrimeric G-protein β subunit controls shoot meristem development and immune responses

**DOI:** 10.1101/797985

**Authors:** Qingyu Wu, Fang Xu, Lei Liu, Si Nian Char, Yezhang Ding, Eric Schmelz, Bing Yang, David Jackson

## Abstract

Heterotrimeric G-proteins are important transducers of receptor signaling, functioning in plants with CLAVATA receptors in control of shoot meristem size, and with pathogen associated molecular pattern (PAMP) receptors in basal immunity. However, whether specific members of the heterotrimeric complex potentiate crosstalk between development and defense, and the extent to which these functions are conserved across species, has not been addressed. Here we used CRISPR/Cas9 to knockout the maize *Gβ* subunit gene, and found that the mutants were lethal, differing from Arabidopsis, where homologous mutants have normal growth and fertility. We show that lethality is not caused by a specific developmental arrest, but by autoimmunity. We used a genetic diversity screen to suppress the lethal *gβ* phenotype, and also identified a new maize *Gβ* allele with weak autoimmune responses but strong development phenotypes. Using these tools, we show that *Gβ* controls meristem size in maize, acting epistatically with *Gα*, suggesting that *Gβ* and *Gα* function in a common signaling complex. Furthermore, we used an association study to show that natural variation in *Gβ* influences maize kernel row number, an important agronomic trait. Our results demonstrate the dual role of *Gβ* in immunity and development in a cereal crop, and suggest that it functions in crosstalk between these competing signaling networks. Therefore, modification of Gβ has the potential to optimize the tradeoff between growth and defense signaling to improve agronomic production.

**Significance:** Cereal crops, such as maize provide our major food and feed. Crop productivity has been significantly improved by selection of favorable architecture and development alleles, however crops are constantly under attack from pathogens, which severely limits yield due to a defense-growth tradeoff. Therefore, it is critical to identify key signaling regulators that control both developmental and immune signaling, to provide basic knowledge to maximize productivity. This work shows that the maize G protein β subunit regulates both meristem development and immune signaling, and suggests that manipulation of this gene has the potential to optimize the tradeoff between yield and disease resistance to improve crop yields.

## Introduction

Shoots are derived from meristems, pool of self-renewing stem cells that initiate new organs from their daughter cells (1). The development of the shoot apical meristem (SAM) is controlled by the CLAVATA (CLV)-WUSCHEL (WUS) feedback signaling pathway (1). This pathway includes a secreted peptide, CLV3, its leucine-rich repeat receptor-like kinase (LRR-RLK) CLV1, and a homeodomain transcription factor WUS, which promotes CLV gene expression and stem cell fate (2–5). CLV1 binds and perceives CLV3 peptide, leading to WUS repression (2–5). A second LRR protein, CLV2, is a receptor-like protein that controls meristem size in parallel to CLV1 (4, 6). The CLV-WUS feedback loop was discovered in the model species Arabidopsis, but is conserved widely, including in cereal crops. Through characterizing maize *fasciated ear* (*fea*) mutants with enlarged inflorescence meristems (IMs), the *THICK TASSEL DWARF1 (TD1), FASCIATED EAR2 (FEA2)* and *ZmCLAVATA3/EMBRYO SURROUNDING REGION-RELATED7* (*ZmCLE7*) genes were identified as orthologs of *CLV1, CLV2*, and *CLV3* respectively (7–11). In addition to the conventional CLV1 receptor, the LRR receptor-like protein FASCIATED EAR3 (FEA3) represses WUS from below, and perceives a distinct CLE peptide, ZmFON2-LIKE CLE PROTEIN1 (ZmFCP1) that is expressed in differentiating cells (10). Therefore, distinct CLV receptors perceive small CLE peptides to maintain the balance of meristem proliferation and differentiation. However, the downstream signaling events from these receptors are not well understood.

Heterotrimeric G-proteins, consisting of Gα, Gβ, and Gγ subunits, transduce signals downstream of receptors (12). In the standard animal model, a GDP-bound Gα associates with a Gβγ dimer and a 7-pass transmembrane (7-TM) G-protein coupled receptor (GPCR) in its inactive state. Upon ligand perception, the GPCR promotes GDP release and binding of GTP by Gα, activating the G-proteins and promoting interaction with downstream effectors (12). However, G-protein signaling in plants appears to be fundamentally different, and whether plants have 7-TM GPCRs is still under debate (13–15). In contrast, emerging evidence suggests that heterotrimeric G-proteins in plants interact with single-pass transmembrane receptors (16–19). For example, the maize G-protein *α* subunit COMPACT PLANT2 (CT2) interacts with the CLV2 ortholog FEA2 to control shoot meristem development, and *ct2* mutants have enlarged SAMs and fasciated ears (16). Similarly, the Arabidopsis G-protein *β* subunit (AGB1) interacts with another CLV-like receptor, RECEPTOR-LIKE PROTEIN KINASE2 (RPK2) to control Arabidopsis SAM development, and Arabidopsis *agb1* mutant SAMs are larger (16, 18).

In addition to their developmental functions, heterotrimeric G-proteins also positively regulate plant immunity. For example, AGB1 and EXTRA-LARGE GTP BINDING PROTEIN2 (XLG2), a non-canonical *Gα* in Arabidopsis, interact with the immune receptor FLAGELLIN SENSITIVE2 (FLS2), as well as with its downstream kinase BOTRYTIS-INDUCED KINASE1 (BIK1), which is stabilized by this interaction (19), thus immunity is compromised in *xlg* or *gβ* mutants (19, 20). RNAi suppression of the rice *Gβ* gene *RGB1* causes browning of internodes and ectopic cell death in roots, phenotypes associated with immune defects (21, 22). However, the functions of monocot G*β* genes in development have never been dissected, because CRISPR/Cas9-derived *rgb1* null mutants die soon after germination (23, 24).

Here, we report that CRISPR/Cas9 induced knockouts of maize *Gβ* (*ZmGB1)* are seedling lethal, distinct from Arabidopsis but similar to rice. We found that lethality was due to autoimmunity, rather than a developmental arrest. We rescued lethality by introgressing *Zmgb1* CRISPR (*Zmgb1*^*CR*^) mutants into a suppressive genetic background, and found that the mutants had larger SAMs and fasciated inflorescences. We also identified a viable allele of *ZmGB1* by map-based cloning of a fasciated ear mutant, *fea*\**183*, which preferentially alleviated immune phenotypes. Our study dissects the dual functions of *Gβ* in shoot meristem development and immune responses, suggesting that modulation of G protein signaling has the potential to optimize the tradeoff between yield and disease resistance in crop plants.

## Results

### Knocking out *ZmGB1* using CRISPR/Cas9 caused lethality due to autoimmunity

Maize Gα (CT2) and Arabidopsis Gα and β subunits control meristem development (16, 18, 25). However, the role of Gβ in meristem regulation in the grasses remains obscure, because rice *Gβ* knockouts are lethal, leading to the proposal that it is essential for growth (23, 24). To study the function of maize G*β*, we used CRISPR/Cas9 to generate multiple alleles, including 1-bp and 136-bp deletions with premature stop codons predicted to result in null alleles (Fig. 1A). Homozygous *Zmgb1*^*CR*^ mutants germinated normally, but arrested and turned yellow then brown, and died at an early seedling stage (Fig. 1B). The necrotic appearance of *Zmgb1*^*CR*^ mutants, along with the known role of *AGB1* in Arabidopsis immune responses (19, 20), prompted us to survey immune markers. We first checked for cell death by staining with trypan blue. *Zmgb1*^*CR*^ mutants were heavily stained compared to wild-type, suggesting that they were undergoing cell death (Fig. 1C). In support of this, 3,3′-diaminobenzidine (DAB) staining showed that H_2_O_2_, another marker for immune responses, accumulated in the mutants (Fig. 1D). We also checked the expression of two immune marker genes, *PATHOGENESIS-RELATED PROTEIN1 (PR1)* and *PR5*, and found that both were significantly higher in *Zmgb1*^*CR*^ mutants (Fig. 1E), as were levels of the defense hormone salicylic acid (Fig. 1F). Similar necrotic phenotypes were found in mutants grown in sterile culture, which together with the up-regulation of immune markers suggests that *Zmgb1*^*CR*^ mutants died because of an autoimmune response.

**Figure 1.**
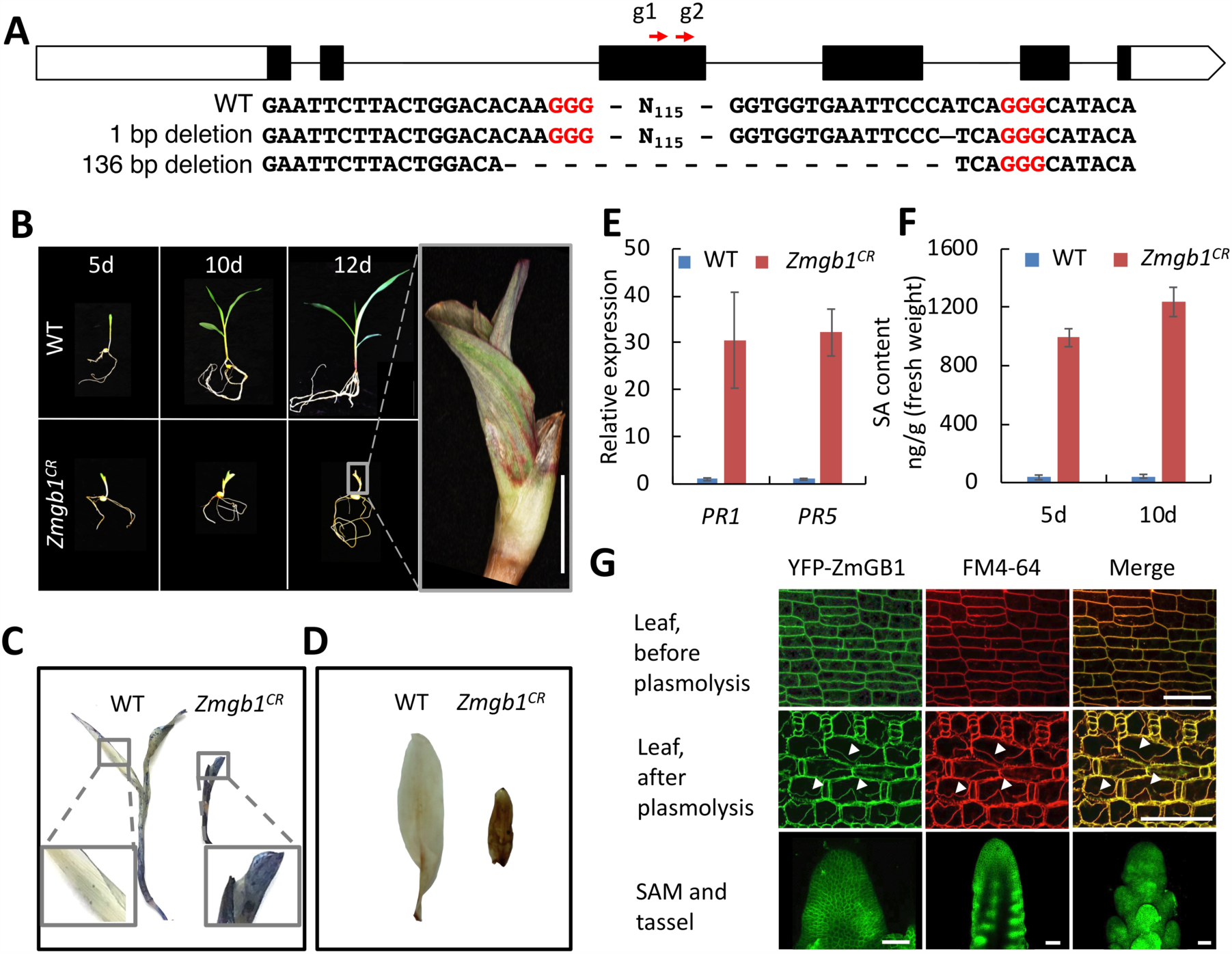
CRISPR/Cas9 knockouts of *ZmGB1* led to autoimmune phenotypes. (A) CRISPR/Cas9 editing of *ZmGB1* produced different frameshift alleles. White boxes indicate 5’ and 3’ UTRs; black boxes indicate exons, and black lines indicate introns. The positions of guide RNAs are indicated by red arrows. (B) *Zmgb1*^*CR*^ mutants were lethal at the seedling stage. The pictures were taken 5, 10, and 12-d post sowing seeds in soil. The upper panels are wild-type (WT) and the lower panels are *Zmgb1*^*CR*^ mutants. Scale bar = 1 cm. Trypan blue (C) and DAB (D) staining of WT and *Zmgb1*^*CR*^ mutants showed increased staining in the mutants. (E) *PR1* and *PR5* expression were up-regulated in the *Zmgb1*^*CR*^ mutants, and both 5-d and 10-d-old *Zmgb1*^*CR*^ mutants accumulated significantly more salicylic acid (SA) (F). For (E and F), *p* value = 0.0001; Student’s t test; N = 3. (G) YFP-SBP-ZmGB1 localizes to membranes in shoot meristems. Upper panels, leaf cells expressing YFP-SBP-ZmGB1 (green), counterstained with FM4-64 (red), both visible as a thin line around the cell. Middle panels, following plasmolysis, YFP-SBP-ZmGB1 (arrows) remained co-localized with FM 4-64. In the lower panels, YFP-SBP-ZmGB1 expression was found through SAM and tassel inflorescence primordia. Scale bars = 50 μm.

To confirm that the phenotypes were due to mutation of *ZmGB1*, and not to an off-target effect of CRISPR/Cas9, we made a translational fusion of the *ZmGB1* genomic sequence with YELLOW FLUORESCENT PROTEIN (YFP)-STREPTAVIDIN-BINDING PEPTIDE (SBP) at its N-terminus, under the control of its native promoter and terminator. This construct was transformed into maize and backcrossed twice to *Zmgb1*^*CR*^ heterozygotes in the B73 background. The YFP-SBP-ZmGB1 transgene was able to complement the lethal phenotypes of *Zmgb1*^*CR*^ mutants (SI Appendix Table S1). Imaging revealed YFP-SBP-ZmGB1 localization to the plasma membrane (Fig. 1G), as expected (26), and confirmed by co-localization with FM4-64 after plasmolysis (Fig. 1G). Consistent with its anticipated role in shoot development, *ZmGB1* was expressed throughout the SAM and inflorescence meristems (Fig. 1G).

Having confirmed the *Zmgb1* phenotypes, we asked why the phenotypes of Arabidopsis *Gβ* mutants (reduced immune response, but overall normal growth and fertility) are weaker than in maize. To ask if this was due to the differences in the Gβ protein, or in immune signaling pathways, we expressed maize *ZmGB1* in Arabidopsis, driven by the native *AGB1* promoter. The *ZmGB1* transgene fully rescued the developmental and immune defects of *agb1* mutants (SI Appendix Fig. S1), suggesting that Gβ function is conserved between maize and Arabidopsis, and that the contrasting immune phenotypes are due to differences in immune signaling pathways.

### *Zmgb1* lethality can be suppressed

The early lethality of *Zmgb1*^*CR*^ plants precluded us from observing their meristem phenotypes. Autoimmune phenotypes are common for proteins that are “guardees”, protected (or guarded) by RESISTANCE (R) proteins (27). Since *R* genes are highly polymorphic across accessions, we attempted to suppress *Zmgb1*^*CR*^ autoimmunity by crossing viable heterozygotes to each of the 25 nested association mapping (NAM) maize diversity lines (28), and screened for suppression in the F2s. Indeed, we found that the lethality of *Zmgb1*^*CR*^ could be partially suppressed after crossing to a tropical maize line, CML103. The suppressed *Zmgb1*^*CR*^ mutants were dwarfed with wider stems, similar to the maize *ct2* (*Gα*) mutants (Fig. 2A) (16), and some of the plants survived to flowering (Fig. 2B). Consistent with this growth recovery, the induction of *PR1* and *PR5* immune marker genes was reduced in the suppressed *Zmgb1*^*CR*^ mutants (SI Appendix Fig. S2), confirming that autoimmunity was also suppressed. We took advantage of these lethality-suppressed *Zmgb1*^*CR*^ mutants to study development of their meristems. The mutants had significantly larger SAMs compared to wild-type sibs (Fig. 2C and D) and fasciated IMs (Fig. 2E), indicating that *ZmGB1* controls both SAM and IM development in maize.

**Figure 2.**
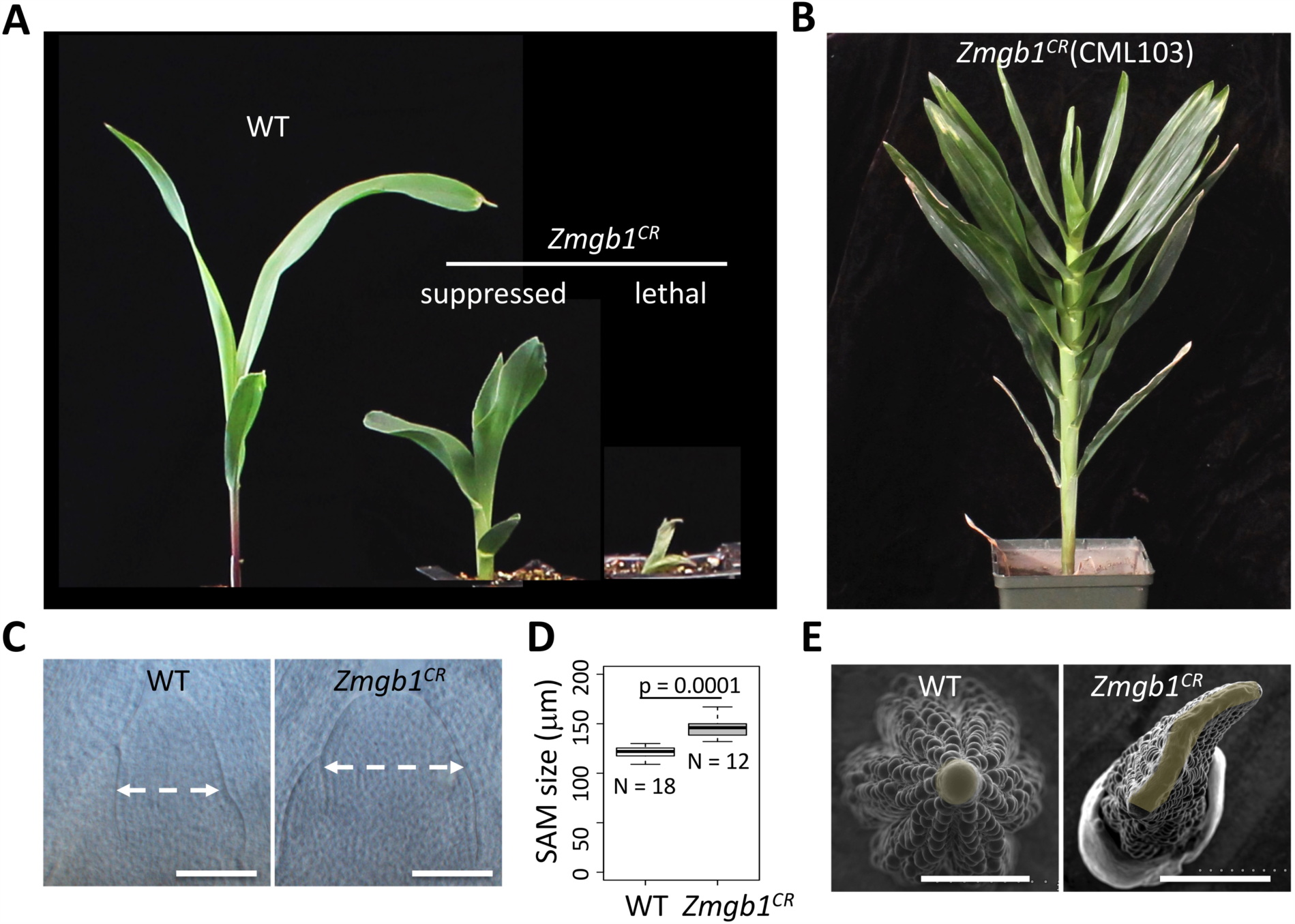
The lethality of *Zmgb1*^*CR*^ mutants was suppressed in the CML103 background. (A) F2 progeny of a cross between *Zmgb1*^*CR*^ heterozygotes and CML103 segregates for lethal and suppressed phenotypes. The pictures are of 7-d-old maize seedlings. (B) The suppressed *Zmgb1*^*CR*^ plants in the CML103 background grew to the adult stage. (C) *Zmgb1*^*CR*^ mutants had enlarged SAMs, quantification in (D) *p* value = 0.0001; Student’s t-test, N=18 for WT and 12 for *Zmgb1*^*CR*^. (E) Top-down view of WT and *Zmgb1*^*CR*^ ear primordia by scanning electron microscopy; IMs are shaded in yellow. Scale bars = 100 μm for (C) and 1 mm for (E).

### A newly identified fasciated ear mutant, *fea*183*, encodes a viable allele of *ZmGB1*

Concurrently, we identified a viable recessive allele of *Zmgb1* by map-based cloning of *fea*183*, a fasciated ear mutant from an ethyl methanesulfonate (EMS)-mutagenesis screen. *fea*183* mutants were semi-dwarfed, and had shorter, wider leaves with prominent lesions (Fig. 3A). They also had striking inflorescence defects, including fasciated ears and compact tassels (Fig. 3B,C and SI Appendix Fig. S3A), reminiscent of *ct2* mutants (16). We analyzed developing ear and tassel primordia using scanning electron microscopy (SEM), and found their inflorescence meristems were significantly enlarged (Fig. 3B, SI Appendix Fig. S3B). In addition to IM defects, *fea*183* mutants had larger shoot apical meristems (Fig. 3D and E). The mutants also had obvious cell death and up-regulation of *PR* genes, suggesting an autoimmune phenotype, albeit much weaker than *Zmgb1*^*CR*^ mutants (SI Appendix Fig. S3C and D). Bulked segregant analysis and map-based cloning delineated the *fea*183* mutation between 257.3MB and 258.9 MB on chromosome 1 (Fig. 3F, SI Appendix Fig. S4A). Whole genome sequencing identified a single non-synonymous mutation within this region, a G-to-A substitution in the fourth exon of *ZmGB1*, leading to a change in the 277^th^ amino acid from aspartic acid to asparagine in one of the WD40 domains (SI Appendix Fig. S4B). This residue is fully conserved across a wide range of species including *S. cerevisiae, C. elegans, H. sapiens*, and Arabidopsis, implying its essential role in Gβ function (SI Appendix Fig. S4C). We next confirmed that *fea*183* encoded an allele of *ZmGB1*, by crossing with *Zmgb1*^*CR*^ heterozygous plants. In the F1, about half of the plants had enlarged IMs and dwarfism, similarly to *fea*183* mutants (Fig. 3G and H), indicating a failure to complement, and that *FEA*183* encodes the maize G protein β subunit. Thus, we renamed *fea*183* as *Zmgb1*^*fea*183*^.

**Figure 3.**
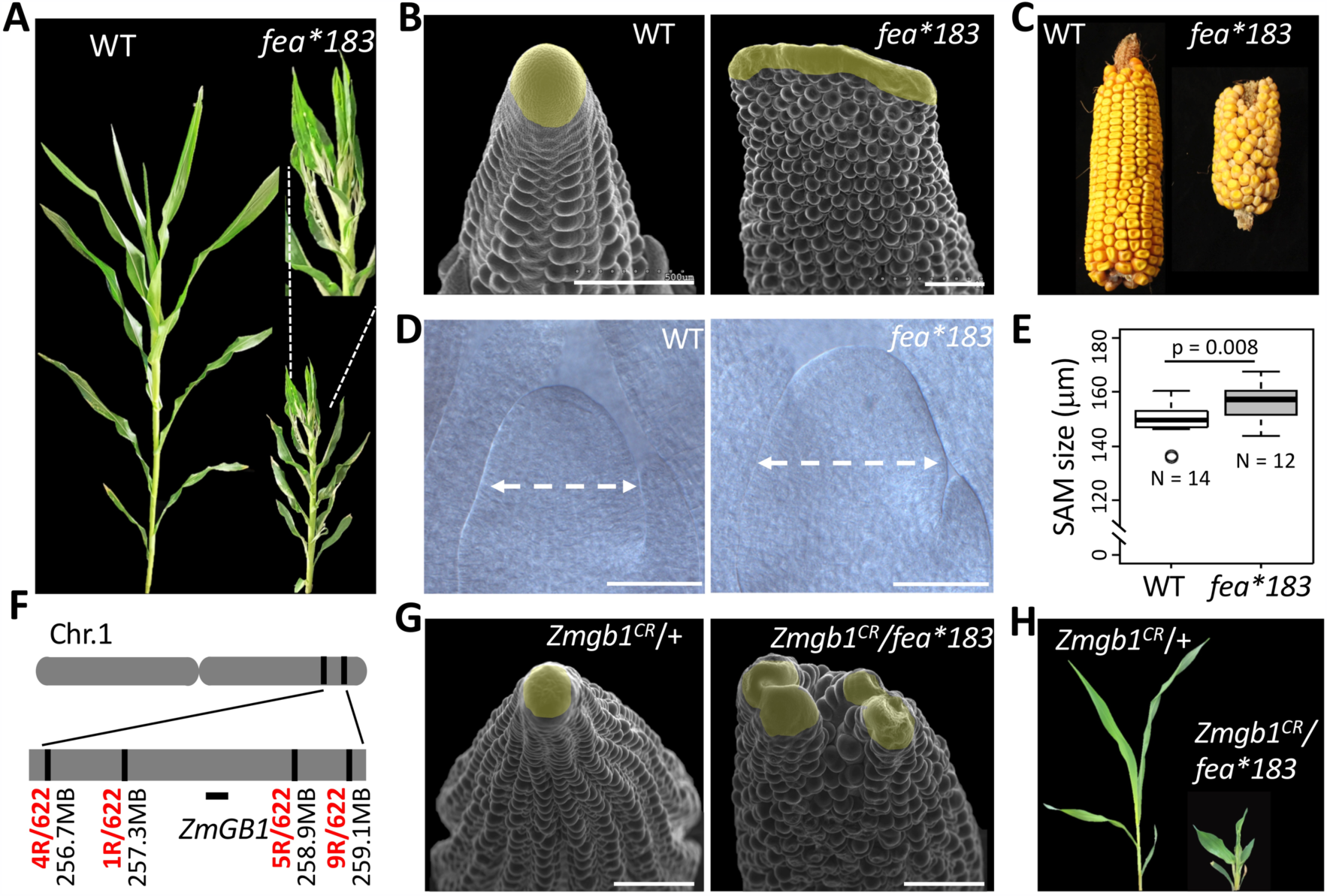
Characterization and mapping of the *fea*183* mutant. (A) *fea*183* plants were semi-dwarfed with upright leaves and lesions (inset dotted lines). (B) SEM images showing that *fea*183* mutant ear primordia had enlarged IMs, shaded in yellow. Scale bars = 500 μm. (C) Representative mature cobs of WT and *fea*183* showing fasciated ear phenotype. (D) Cleared SAM images of 12-d-old WT and *fea*183* mutants. Scale bars = 100 μm. (E) *Zmgb1*^*CR*^ mutants had bigger SAMs; *p* value = 0.008, Student’s t-test, N = 14 for WT and 12 for *fea*183.* (F) Positional cloning of *fea*183* mutants identified *ZmGB1* as the candidate gene. The vertical lines indicate the position of markers used. The number of recombinants at each position is listed in red. (G) *fea*183* failed to complement *Zmgb1*^*CR*^ in IM development; SEM images of ear primordia. Scale bars = 500 μm. (H) *fea*183* failed to complement *Zmgb1*^*CR*^ seedling development, pictures are of 2-week-old seedlings.

We next asked how the D^277^N mutation affected *Zmgb1*^*fea*183*^ function, by comparing it with human Gβ, HsGB1, and guided by a structure of the human G protein complex (29). The D^277^ residue in Zmgb1^fea*183^ aligned to D^254^ in HsGB1 (29) (Fig. 4A), which lies at the interface of Gβ and Gγ (Fig. 4B). We thus asked whether the this residue was required to form the heterotrimeric complex, using a yeast-3-hybrid (Y3H) experiment (30). We found that unlike the wild-type protein, the Zmgb1^fea*183^ protein could not form a complex with a maize Gγ subunit (ZmRGG2) and Gα/CT2, or with any of the XLG proteins (Fig. 4C), indicating that Zmgb1^fea*183^ was unable to form a heterotrimeric complex, and suggesting it resembles a null allele.

**Figure 4.**
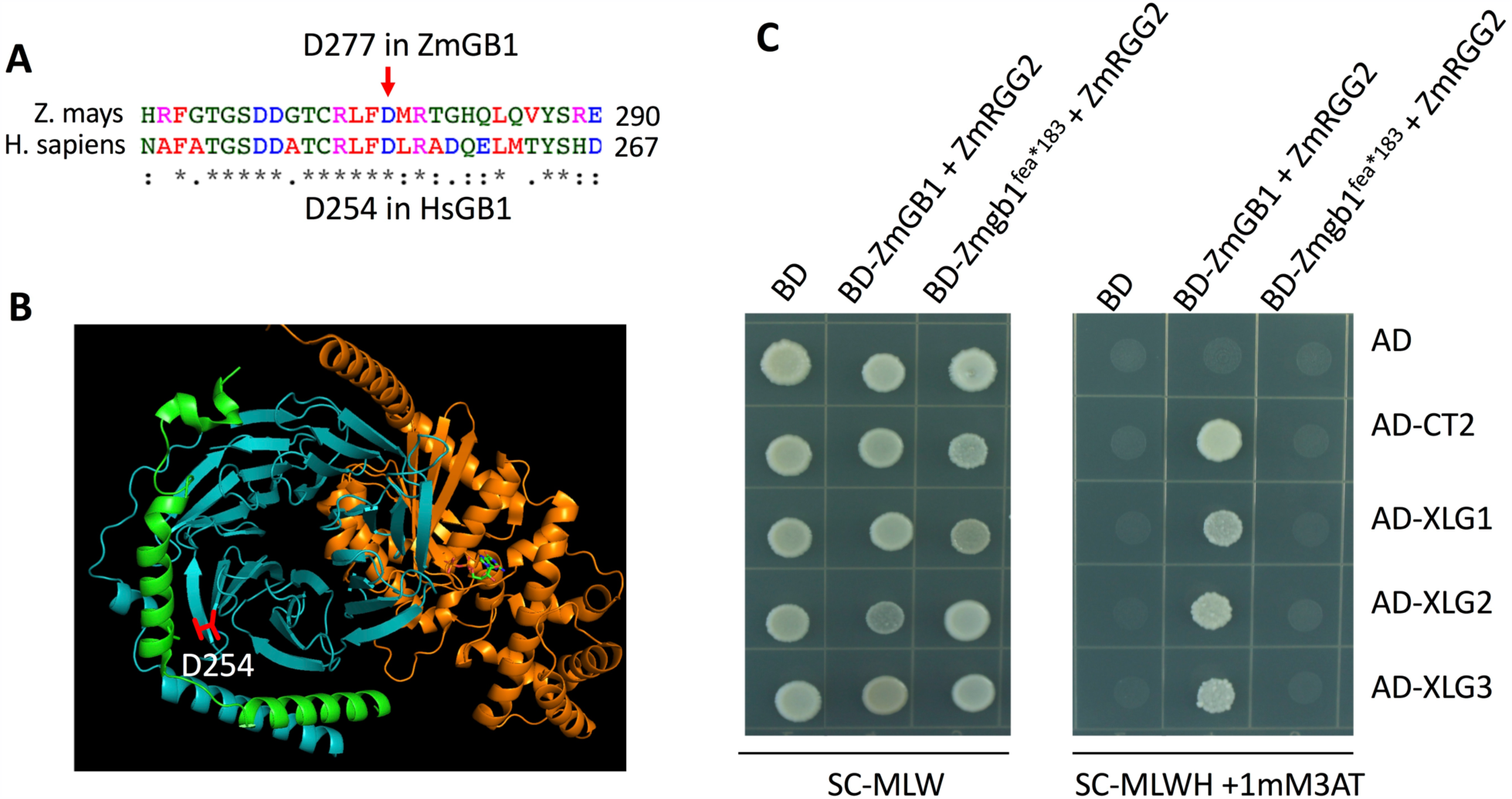
*Zmgb1*^*fea*183*^ failed to form a protein complex with G*α* and Gγ subunits. (A) The D277 residue mutated in *Zmgb1*^*fea*183*^ aligns to D254 in Human HsGB1. (B) D254 highlighted in red in HsGB1 is located at the Gβ and Gγ interface. Viewed by PyMoL, with Gα subunit in orange, Gβ subunit in cyan and Gγ subunit in green. (C) ZmGB1 and the ZmRGG2 Gγ subunit formed complexes with Gα/ CT2 or XLGs in a Y3H assay, while Zmgb1^fea*183^ could not. ZmGB1 was fused with the BD domain and co-expressed with RGG2 using a pBridge construct (Clontech). Gα/ CT2 or individual XLG proteins were fused with the AD domain in the pGADT7 vector. Yeast growth on synthetic complete -Met -Trp -Leu (SC-MLW) medium confirmed transformation and cell viability. Interactions were assayed on SC-Met -Trp -Leu -His (SC-MLWH) medium supplemented with 1 mM 3-AT.

### ZmGB1 functions in the CLAVATA pathway

To further decipher the role of ZmGB1, we made double mutants using the *Zmgb1*^*fea*183*^ allele with other meristem regulatory genes, including *fea2, ct2*, and *fea3* (8, 10, 16), and measured meristem size in segregating populations. The SAMs and ear inflorescence meristems of *Zmgb1;fea2* double mutants were not obviously different from those of *fea2* single mutant, indicating that *fea2* is epistatic to *Zmgb1*, and suggesting that they act in a common pathway (Fig. 5A,B,C). Similarly, IMs of *Zmgb1;ct2* double mutants were no more fasciated than either single mutant, suggesting that ZmGB1 and CT2/Gα function together in regulating IM development (Fig. 5D). However, vegetative SAMs of *Zmgb1;ct2* double mutants were more severely affected than the single mutant, presumably because CT2 acts redundantly with ZmXLGs during vegetative development (30) (Fig. 5E,F). Finally, *Zmgb1;fea3* double mutants had significantly larger SAMs and more strongly fasciated IMs than either single mutant (Fig. 5G,H,I), indicating an additive genetic effect and that *Zmgb1* and *fea3* act in different pathways in both SAM and IM regulation, in line with previous observations (31). In summary, our data suggest that ZmGB1 functions together with CT2/Gα in inflorescence development, downstream of the FEA2 CLAVATA receptor.

**Figure 5.**
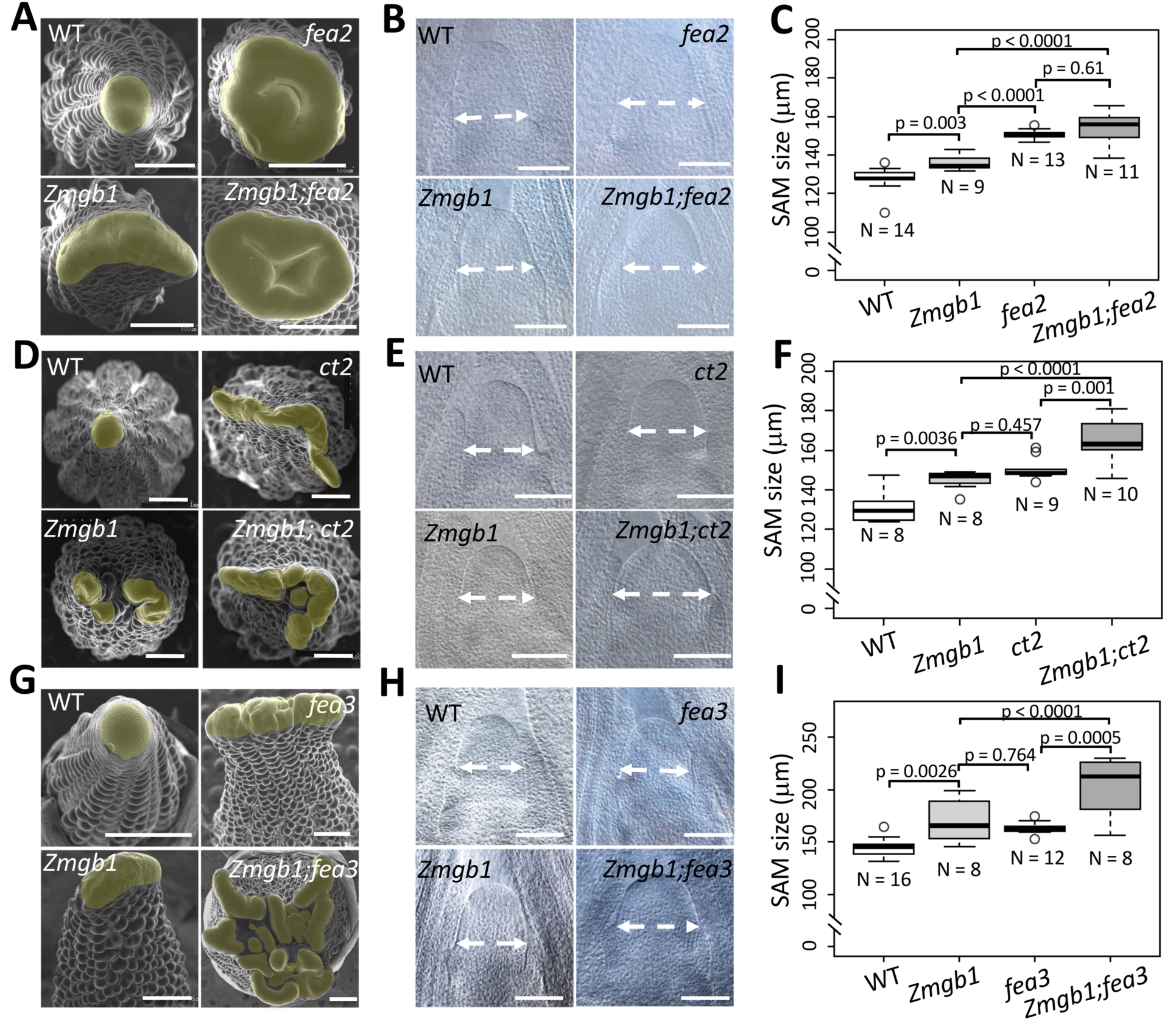
ZmGB1 functions in a CLAVATA pathway. (A) SEM images of WT, *Zmgb1, fea2* and *Zmgb1;fea2* ear primordia. The double mutants showed similar IMs as the *fea2* single mutant. (B) Representative SAM pictures from 16d-old WT, *Zmgb1, fea2* and *Zmgb1;fea2* plants. (C) SAM size quantification showed that the SAM size of the *Zmgb1;fea2* double mutants was indistinguishable from *fea2* single mutants. (D) SEM images of WT, *Zmgb1, ct2* and *Zmgb1;ct2* ear primordia. The double mutants showed similar IMs as the single mutants. (E) Representative SAM pictures of WT, *Zmgb1, ct2* and *Zmgb1;ct2* plants. (F) The SAM size of the *Zmgb1;ct2* double mutant was significantly increased compared to single mutants. (G) SEM images of WT, *Zmgb1, fea3* and *Zmgb1;fea3* ear primordia. The double mutants showed significantly enlarged IMs compared to the single mutants. (H) Representative SAM pictures of 16d-old WT, *Zmgb1, fea3* and *Zmgb1;fea3* plants. (I) SAM size of the *Zmgb1;fea3* double mutants was significantly larger than the single mutants. ANOVA analysis was performed using the R program for (C, F, I). The *p* values, mean values and replicate numbers are indicated in the figures. Scale bar = 500 μm for (A, D, G) and = 100 μm for (B, E, H).

### *ZmGB1* associates with maize kernel row number (KRN)

KRN is an important agronomic trait that directly contributes to yield (10, 32, 33). Natural or induced variation in *FEA2* or *FEA3* is associated with KRN, and manipulation of *CT2* also enhances KRN (10, 30, 32). We therefore asked if *ZmGB1* also associates with this yield trait by conducting a candidate gene association study using a maize association panel of 368 diverse inbred lines (34). Indeed, we found that five SNPs in the first and third exons of *ZmGB1* significantly associate with maize KRN (Fig. 6A). These five KRN associated SNPs can form four kinds of haplotypes among the 368 lines and two of them (Hap3 and Hap4) have significantly more kernel rows than the other two (Fig. 6B). For example, the Hap4 has 1.5 to 2.5 more kernel rows comparing with Hap2 and Hap1, respectively (Fig. 6B). However, the frequencies of favorable Hap3 and Hap4 in the association panel are only 2.17% and 4.07%, implying that the favorable *ZmGB1* alleles haven’t been selected during maize breeding. Therefore, our results suggesting that natural variation in *ZmGB1* influences IM size and KRN, with the potential to benefit maize yields.

**Figure 6.**
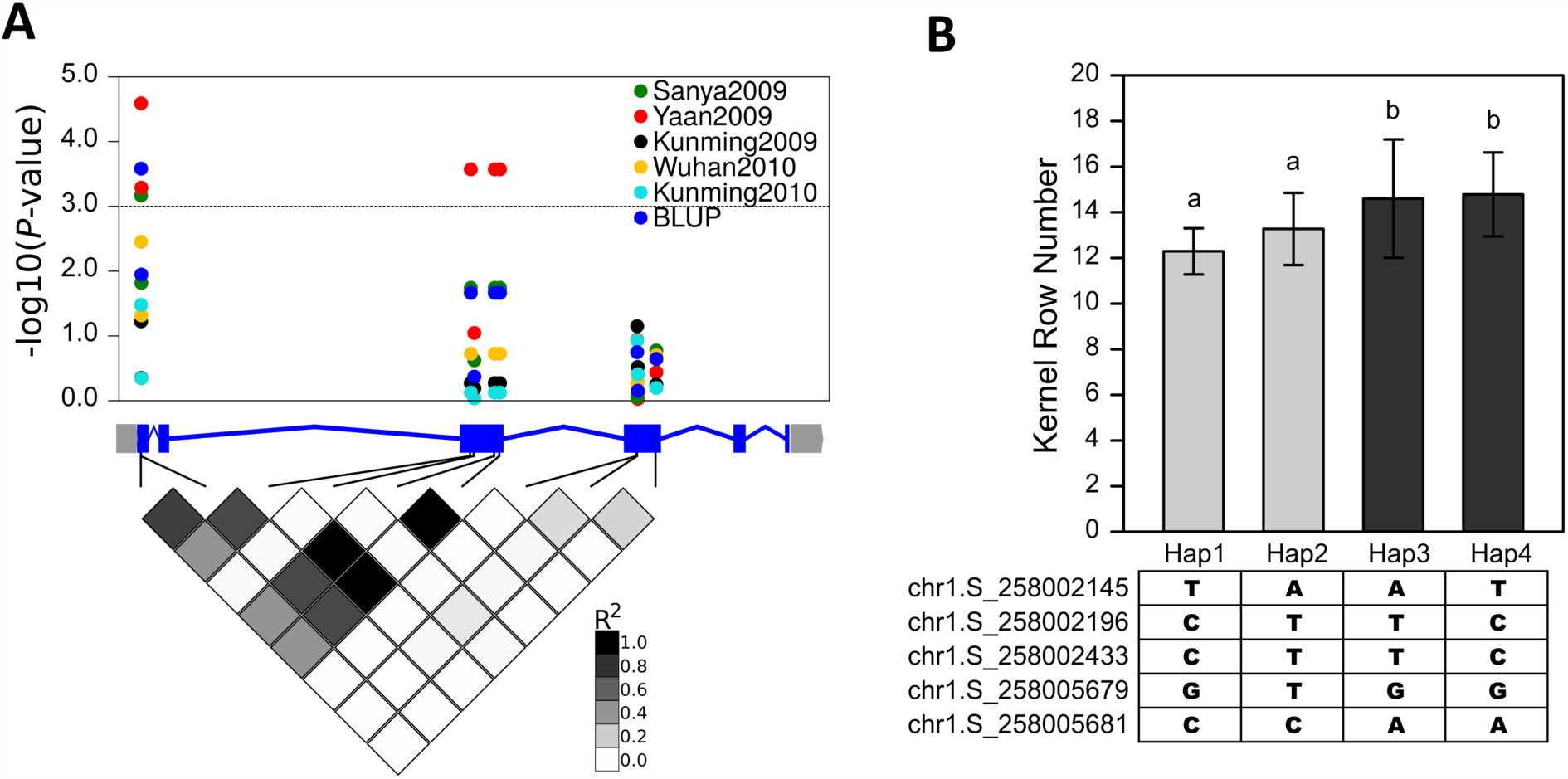
The association analysis of *ZmGB1* with kernel row number. (A) The dots show multiple coding SNPs that associate positively with KRN over multiple environments, and their BLUP (best linear unbiased prediction) data. 368 diverse inbred lines were used in the association analysis using the MLM + Q model. Shaded diamonds below gene model show the SNP linkage disequilibrium (LD) by pairwise R^2^. (B) The haplotype analysis using the five KRN associated SNPs and the KRN (BLUP) of these haplotypes in the association panel, multiple comparisons *P*-Value < 0.05.

## Discussion

Heterotrimeric G-proteins are important signal transducers that control many biological processes across a wide range of species (12, 35). They also control many important agronomic traits in cereals (16, 23, 30, 36-39), and understanding G-protein signaling requires a study of each subunit. Rice *Gβ* CRISPR null mutants undergo an early developmental arrest and death, but the underlying mechanism was unclear (23, 24). Here, we show that maize *Gβ* null alleles are also lethal, and this is due to autoimmunity, not to specific developmental defects. We suppressed the *Gβ* lethal phenotype in the CML103 tropical maize genetic background, and identified a viable EMS allele, allowing developmental analysis *Gβ* meristems. Using the suppressed CRISPR null and the viable *Zmgb1*^*fea*183*^ alleles, we show that G*β* controls shoot meristem development. Our results suggest that G*β* interacts with different downstream effectors to independently function in immune and development signaling.

An important question is why only monocot *Gβ* mutants, such as in rice or maize, but not Arabidopsis mutants, develop autoimmunity? Intriguingly, the Arabidopsis *Gβ* mutant *agb1* has reduced immune response, contrasting with the autoimmune phenotype in rice or maize (19, 20). Expression of maize *Gβ* fully complemented the immune defects of Arabidopsis *agb1* mutants (SI Appendix Fig. S1), suggesting that G*β* protein function is conserved, and that the contrasting phenotypes are due to differences in immune signaling pathways. Plants have a two-tiered immune system. First, Pathogen-Associated Molecular Pattern (PAMP) receptors recognize conserved microbial elicitors and induce Pattern Triggered Immunity (PTI) (40–42). To overcome PTI, pathogens have evolved effectors that they secrete into plant cells to interfere with PAMP signaling, and in turn, plants evolved *R* genes to activate the stronger “Effector Triggered Immunity” (ETI), which often results in programmed cell death (43–46). Some R proteins guard native plant proteins, known as ‘guardees’ that are targeted by pathogen effectors. Thus, mutation of a ‘guardee’ may mimic the presence of a pathogen, and activate the guarding R protein, resulting in an autoimmune phenotype (47). It is therefore reasonable to speculate that grass Gβ proteins function as immune guardees. Supporting this hypothesis, Gβ has 7 WD-40 domains, and forms a propeller structure, similar to some other effector targets (48). Our hypothesis explains why the immune phenotypes of *Zmgb1*^*fea*183*^ mutants are weaker, because presumably that allele accumulates some, albeit mutant, Gβ protein. *R* genes are highly polymorphic across accessions, and our results suggest that Gβ is guarded in monocots rice and maize, but not Arabidopsis (48). To test this hypothesis, further studies are needed to identify the gene(s) that are responsible for the suppression of *Zmgb1* lethality in CML103.

Our genetic analyses suggest that ZmGB1 works in a common pathway with FEA2 and CT2/Gα, but independent from FEA3. *fea2* was epistatic to both *ct2 (Gα)* and *Zmgb1* in inflorescence meristem fasciation, suggesting that both G protein subunits function together downstream of the FEA2 receptor. However, *ct2/Gα* and *Zmgb1* phenotypes were additive in the SAM, which could be explained by redundancy with the non-canonical Gα proteins, or XLGs, in the SAM (30). However, *Zmxlg* triple mutants are also lethal (30), preventing us from making higher order mutants in maize. Identification of a viable genetic background for *Gα* higher order mutants would help to address this question.

Geneticists and breeders have used QTL and genome wide association (GWA) analysis to identify genes involved in yield traits. Several yield QTL that correspond to heterotrimeric G proteins or CLV-WUS genes have been cloned in rice, maize, and tomato (33, 37, 49-51). For example, *FEA2* is a QTL responsible for variation in maize kernel row number (KRN) (32), and one of the rice *Gγ* genes, *GS3* is a QTL for grain length, weight, and thickness (39), while another rice *Gγ* gene, *DEP1*, is a QTL for rice grain yield and nitrogen use efficiency (37, 38). These studies indicate that G proteins and other meristem regulators have the potential to benefit yield traits. In this study, we found that *ZmGB1* also associated significantly with KRN under multiple environments, suggesting that it also contribute to quantitative variation in KRN, and engineering of weak *ZmGB1* alleles may be an effective approach to improve yield traits (10, 32, 52).

Improvement of crop productivity involved selection of favorable architecture and development alleles. Despite these striking innovations, crops are constantly under attack from pathogens. However, turning on defense signaling often causes a reduction in growth and yield (53, 54). This defense-growth tradeoff results from defense signaling being intertwined with physiological networks regulating plant fitness (53). It is therefore critical to understand developmental and immune signaling crosstalk to provide basic knowledge to maximize productivity. Our study shows that *ZmGB1* is a critical regulator in both meristem development and immunity; therefore, this gene has the potential to optimize defense-development tradeoffs to improve agronomic production.

## Materials and Methods

The *Zmgb1*^*CR*^ alleles were created using CRISPR/Cas9, and the *Zmgb1*^*fea*183*^ allele was from an EMS mutagenesis screen by Gerry Neuffer. Complete details regarding materials, experimental methods, and data analyses are provided in the SI Appendix.

## Acknowledgements

We thank Gerry Neuffer for generating the *Zmgb1*^*fea*183*^ EMS stocks. We acknowledge funding from the Agriculture and Food Research Initiative competitive grant no. 2017-06299 and 2015-06319 of the USDA National Institute of Food and Agriculture to D.J., and from the National Science Foundation grant ISO-1936492 to B.Y.. Q.W. was supported by the National Science and Technology Major Project for Development of Transgenic Organisms (2019ZX08010004-002) and Innovation Program of Chinese Academy of Agricultural Sciences. F.X. was supported by Human Frontier Science Program Long-Term Fellowship LT000227/2016.

## References

1. Wu QY, Xu F, & Jackson D (2018) All together now, a magical mystery tour of the maize shoot meristem. Curr Opin Plant Biol 45:26–35.

2. Clark SE, Williams RW, & Meyerowitz EM (1997) The CLAVATA1 gene encodes a putative receptor kinase that controls shoot and floral meristem size in Arabidopsis. Cell 89(4):575–585.

3. Fletcher JC, Brand U, Running MP, Simon R, & Meyerowitz EM (1999) Signaling of cell fate decisions by CLAVATA3 in Arabidopsis shoot meristems. Science 283(5409):1911–1914.

4. Jeong S, Trotochaud AE, & Clark SE (1999) The Arabidopsis CLAVATA2 gene encodes a receptor-like protein required for the stability of the CLAVATA1 receptor-like kinase. The Plant cell 11(10):1925–1934.

5. Schoof H, et al. (2000) The stem cell population of Arabidopsis shoot meristems is maintained by a regulatory loop between the CLAVATA and WUSCHEL genes. Cell 100(6):635–644.

6. Muller R, Bleckmann A, & Simon R (2008) The receptor kinase CORYNE of Arabidopsis transmits the stem cell-limiting signal CLAVATA3 independently of CLAVATA1. The Plant Cell 20(4):934–946.

7. Taguchi-Shiobara F, Yuan Z, Hake S, & Jackson D (2001) The FASCIATED EAR2 gene encodes a leucine-rich repeat receptor-like protein that regulates shoot meristem proliferation in maize. Genes Dev 15(20):2755–2766.

8. Bommert P, et al. (2005) THICK TASSEL DWARF1 encodes a putative maize ortholog of the Arabidopsis CLAVATA1 leucine-rich repeat receptor-like kinase. Development 132(6):1235–1245.

9. Goad DM, Zhu C, & Kellogg EA (2016) Comprehensive identification and clustering of CLV3/ESR-related (CLE) genes in plants finds groups with potentially shared function. New Phytol 216(2):605–616.

10. Je B, et al. (2016) Signaling from maize organ primordia via FASCIATED EAR3 regulates stem cell proliferation and yield traits. Nature Genetics 48(7):785–791.

11. Rodriguez-Leal D, et al. (2019) Evolution of buffering in a genetic circuit controlling plant stem cell proliferation. Nat Genet 51(5):786–792.

12. Gomes I, et al. (2016) G Protein-Coupled Receptor Heteromers. Annual Review of Pharmacology and Toxicology 56:403–425.

13. Jones JC, Temple BRS, Jones AM, & Dohlman HG (2011) Functional reconstitution of an atypical G protein heterotrimer and regulator of G protein signaling protein (RGS1) from Arabidopsis thaliana. The Journal of biological chemistry 286(15):13143–13150.

14. Urano D & Jones AM (2013) Round up the usual suspects; A Comment on Nonexistent Plant GPCRs. Plant Physiol. 161(3):1097–1102.

15. Taddese B, et al. (2014) Do Plants Contain G Protein-Coupled Receptors? Plant Physiol 164(1):287–307.

16. Bommert P, Je BI, Goldshmidt A, & Jackson D (2013) The maize Galpha gene COMPACT PLANT2 functions in CLAVATA signalling to control shoot meristem size. Nature 502(7472):555–558.

17. Choudhury SR & Pandey S (2013) Specific Subunits of Heterotrimeric G Proteins Play Important Roles during Nodulation in Soybean. Plant Physiol 162(1):522–533.

18. Ishida T, et al. (2014) Heterotrimeric G proteins control stem cell proliferation through CLAVATA signaling in Arabidopsis. Embo Reports 15(11):1202–1209.

19. Liang XX, et al. (2016) Arabidopsis heterotrimeric G proteins regulate immunity by directly coupling to the FLS2 receptor. Elife 5:e13568.

20. Liu JM, et al. (2013) Heterotrimeric G Proteins Serve as a Converging Point in Plant Defense Signaling Activated by Multiple Receptor-Like Kinases. Plant Physiol 161(4):2146–2158.

21. Utsunomiya Y, et al. (2011) Suppression of the rice heterotrimeric G protein beta-subunit gene, RGB1, causes dwarfism and browning of internodes and lamina joint regions. Plant J 67(5):907–916.

22. Urano D, Leong R, Wu T, & Jones AM (2019) Quantitative morphological phenomics of rice G protein mutants portend autoimmunity. Developmental Biology:doi.org/10.1016/j.ydbio.2019.1009.1007.

23. Sun SY, et al. (2018) A G-protein pathway determines grain size in rice. Nat Commun 9.

24. Gao Y, et al. (2019) The heterotrimeric G protein beta subunit RGB1 is required for seedling formation in rice. Rice 12.

25. Urano D, et al. (2016) Saltational evolution of the heterotrimeric G protein signaling mechanisms in the plant kingdom. Science Signaling 9:ra93.

26. Kato C, et al. (2004) Characterization of heterotrimeric G protein complexes in rice plasma membrane. Plant J 38(2):320–331.

27. van der Hoorn RA & Kamoun S (2008) From Guard to Decoy: a new model for perception of plant pathogen effectors. Plant Cell 20(8):2009–2017.

28. McMullen MD, et al. (2009) Genetic Properties of the Maize Nested Association Mapping Population. Science 325(5941):737–740.

29. Maeda S, et al. (2018) Development of an antibody fragment that stabilizes GPCR/G-protein complexes. Nat Commun 9.

30. Wu Q, Regan M, Furukawa H, & Jackson D (2018) Role of heterotrimeric Galpha proteins in maize development and enhancement of agronomic traits. PLoS Genet 14(4):e1007374.

31. Je B, et al. (2018) The CLAVATA receptor FASCIATED EAR2 responds to distinct CLE peptides by signaling through two downstream effectors. Elife 7:e35673.

32. Bommert P, Nagasawa NS, & Jackson D (2013) Quantitative variation in maize kernel row number is controlled by the FASCIATED EAR2 locus. Nature Genetics 45(3):334–337.

33. Liu L, et al. (2015) KRN4 Controls Quantitative Variation in Maize Kernel Row Number. Plos Genetics 11:e1005670.

34. Yang XH, et al. (2011) Characterization of a global germplasm collection and its potential utilization for analysis of complex quantitative traits in maize. Mol Breeding 28(4):511–526.

35. Urano D & Jones AM (2014) Heterotrimeric G Protein-Coupled Signaling in Plants. Annual Review of Plant Biology 65:365–384.

36. Liu Q, et al. (2018) G-protein beta gamma subunits determine grain size through interaction with MADS-domain transcription factors in rice. Nat Commun 9.

37. Huang X, et al. (2009) Natural variation at the DEP1 locus enhances grain yield in rice. Nature genetics 41(4):494–497.

38. Sun H, et al. (2014) Heterotrimeric G proteins regulate nitrogen-use efficiency in rice. Nat Genet 46(6):652–656.

39. Fan C, et al. (2006) GS3, a major QTL for grain length and weight and minor QTL for grain width and thickness in rice, encodes a putative transmembrane protein. Theoretical and applied genetics. 112(6):1164–1171.

40. Boller T & Felix G (2009) A Renaissance of Elicitors: Perception of Microbe-Associated Molecular Patterns and Danger Signals by Pattern-Recognition Receptors. Annual Review of Plant Biology 60:379–406.

41. Gómez-Gómez L & Boller T (2000) FLS2: an LRR receptor-like kinase involved in the perception of the bacterial elicitor flagellin in Arabidopsis. Molecular cell 5(6):1003–1011.

42. Monaghan J & Zipfel C (2012) Plant pattern recognition receptor complexes at the plasma membrane. Curr Opin Plant Biol 15(4):349–357.

43. Bent AF & Mackey D (2007) Elicitors, effectors, and R genes: The new paradigm and a lifetime supply of questions. Annu Rev Phytopathol 45:399–436.

44. Caplan JL, Mamillapalli P, Burch-Smith TM, Czymmek K, & Dinesh-Kumar SP (2008) Chloroplastic protein NRIP1 mediates innate immune receptor recognition of a viral effector. Cell 132(3):449–462.

45. Chisholm ST, Coaker G, Day B, & Staskawicz BJ (2006) Host-microbe interactions: Shaping the evolution of the plant immune response. Cell 124(4):803–814.

46. Jones JD & Dangl JL (2006) The plant immune system. Nature 444(7117):323–329.

47. Chakraborty J, Ghosh P, & Das S (2018) Autoimmunity in plants. Planta 248(4):751–767.

48. Sarris PF, Cevik V, Dagdas G, Jones JDG, & Krasileva KV (2016) Comparative analysis of plant immune receptor architectures uncovers host proteins likely targeted by pathogens. Bmc Biol 14.

49. Jiao Y, et al. (2010) Regulation of OsSPL14 by OsmiR156 defines ideal plant architecture in rice. Nature genetics 42(6):541–544.

50. Liu J, et al. (2017) GW5 acts in the brassinosteroid signalling pathway to regulate grain width and weight in rice. Nat Plants 3:17043.

51. Miura K, et al. (2010) OsSPL14 promotes panicle branching and higher grain productivity in rice. Nature genetics 42(6):545–549.

52. Rodriguez-Leal D, Lemmon ZH, Man J, Bartlett ME, & Lippman ZB (2017) Engineering Quantitative Trait Variation for Crop Improvement by Genome Editing. Cell 171(2):470–480.

53. Karasov TL, Chae E, Herman JJ, & Bergelson J (2017) Mechanisms to Mitigate the Trade-Off between Growth and Defense. Plant Cell 29(4):666–680.

54. Huot B, Yao J, Montgomery BL, & He SY (2014) Growth-defense tradeoffs in plants: a balancing act to optimize fitness. Mol Plant 7(8):1267–1287.

