## Supporting information for "The maize heterotrimeric G-protein β subunit controls shoot meristem development and immune responses"

### Materials and Methods

#### Plant growth condition and map-based cloning

Maize plants were grown at the Uplands Farm Agricultural Station at Cold Spring Harbor, New York between June and October, or in the greenhouse. The light cycle in the greenhouse was 16/8 h light/dark and the temperature was between 26 - 28 $^{o}$C.

The *fea*183* mutant was obtained from the Maize Genetics Stock Center, and was originally isolated as a fasciated ear mutant from an ethyl methanesulfonate (EMS)-mutagenized screen in A619 inbred maize (original pedigree 03Il-A619-TR183). *fea*183* mutants were outcrossed to B73 inbreds and the F1 plants were then self-crossed to create a segregating F2 population for mapping. Pooled DNAs from ~30 mutants or the same number of normal ear plants from the segregating F2 population were used for bulked segregant analysis (BSA) using a maize SNP50 chip (Illumina, Inc.). The BSA analysis revealed a strong linkage of the mutation on Chromosome 1 between 242-270 Mb. Using ~350 F2 mutants, we further fine-mapped the mutation to a 1.6 Mbp region containing approximately 70 genes. Whole genome sequencing (Illumina, ~ 20x coverage) using pooled DNAs from ~50 fasciated ear mutants from the F2 family was carried out and GATK 3.5 package was used to call SNPs. After filtering Hapmap SNPs, only one non-synonymous mutation was found within the mapped region, and was predicted to cause an amino acid change in ZmGB1.

#### Knockout of *ZmGB1* using CRISPR/Cas9

CRISPR/Cas9 was used to create *ZmGB1* knockouts in maize genotype Hi-II through *Agrobacterium*-mediated transformation (1). Guide RNA design was based on the B73 reference genome sequence as described (2). Two guide RNAs (gRNAs) were designed to target the coding region of *ZmGB1*. One pair of oligonucleotides were synthesized and annealed to construct the sequence of gRNA for one target site. A second gRNA was constructed sequentially in pENTR-gRNA1, and the resulting gRNA cassette, was mobilized into pGW-Cas9 through Gateway recombination. The CRISPR/Cas9 construct was transferred into the *Agrobacterium* strain EHA101, then into maize HiII via transformation. 51 plants from 8 transgenic events were obtained, and the genomic regions surrounding the target sites were PCR- amplified and Sanger sequenced. The trace files (chromatograms) were manually analyzed to determine the genotypes of individual CRISPR plants. Oligonucleotide information is provided in Table S2.

**SAM measurements**

Maize seedlings were grown in the greenhouse for indicated days then dissected and fixed in FAA (10% formalin, 5% acetic acid, and 45% ethanol). The tissue clearing and SAM measurements were performed as described (3) using a Leica DMRB microscope with a Leica Micropublisher 5.9 RTV digital camera system. The SAM size was measured using Image J.

**qRT-PCR**

8d-old shoot tissues were harvested for measuring *PR1* (GRMZM2G465226) and *PR5* (GRAMZM2G402631) expression. Total RNA was extracted using the Direct-zol RNA extraction kit (Zymo Research). cDNA was synthesized using the iScript reverse transcription supermix (Bio-Rad) according to the manufacturer's manual. qRT-PCR was performed on a CFX96Real-Time system (Bio-Rad). The relative expression level of the targeted gene was normalized using *ZmUBIQUITIN* (3). Primers are listed in Table S2.

**Trypan blue staining**

Staining was performed using 10d-old wild-type and *Zmgb1* mutants, as described with slight modifications (4). Briefly, the shoot was immersed in lactophenol containing 2.5 mg/ml trypan blue, and heated in boiling water for 1 min, then cooled and kept at room temperature overnight. The tissues were cleared in chloral hydrate solution (25 g chloral hydrate in 10 ml $H_2O$) for 24 h at room temperature.

**DAB staining**

Leaves from 10-d-old wild-type and *Zmgb1* mutants were immersed in 1mg/mL DAB at room temperate in the dark for 4 h, then cleared by boiling in 95% ethanol for 10 min. $H_2O_2$ accumulation was detected as brown stain after DAB staining.

**Scanning electron microscopy**

Scanning electron microscopy was performed on fresh ears or tassels using a Hitachi S-3500N SEM.

**Yeast-3-hybrid experiment**

The yeast-3-hybrid experiment was carried out as described in (3). The pBridge-BD-Zmgb1$^{fea*183}$-RGG2 construct was made by site-directed mutagenesis PCR using pBridge-BD-ZmGB1-RGG2 as the template with the primers listed in Table S2.

***ZmGB1* transgene construct and imaging**

*YFP-SBP-ZmGB1* was constructed by amplification of genomic fragments and fusing with the YFP-SBP tag in-frame at the N-terminus using the MultiSite Gateway Pro system (Invitrogen) according to the manufacturer's manual. All fragments were amplified using KOD Xtreme hot start polymerase (Millipore Sigma). The entry clones were assembled in the pTF101 Gateway compatible binary vector and introduced into the EHA101 Agrobacterium strain, and transformed into maize Hi-II background by the Iowa State University Plant Transformation Facility. The T0 plants were backcrossed twice with *Zmgb1*$^{CR}$/+ in the B73 background.

For confocal microscopy, tissues were dissected and counterstained with 1 mg/ml FM4-64 solution (Molecular Probes) in water for 5 min. Images were taken with a Zeiss LSM 710 microscope, using 514 nm laser excitation and 520-560 nm emission for detection of the YFP and 585-750 nm laser excitation for detection of FM4-64. For plasmolysis, leaf tissues were incubated in 30% glycerol for 10 min.

**Arabidopsis *agb1* mutant complementation**

The CDS of *ZmGB1* was driven by a 791 bp promoter region of the Arabidopsis *AGB1* gene, and a 665 bp *AGB1* 3' UTR region cloned into the pTF101 Gateway compatible binary vector using the MultiSite Gateway Pro system (Invitrogen), as described (5). The construct was introduced into the GV3101 Agrobacterium strain then transformed into plants carrying the *agb1-2* allele in Col-0 background. Oxidative burst assays were performed as described with slight modification (6). Briefly, leaf strips from 5-week old soil-grown Arabidopsis plants grown under short day conditions (8h light/16h dark) were incubated in water for 12 h in a 96-well plate before treating with luminescence detection buffer (20 μM luminol and 10 mg/ml horseradish peroxidase) containing 1 μM flg22. Luminescence was recorded using a BioTek Synergy H4 fluorescence microplate reader.

**Quantification of endogenous salicylic acid (SA) level by GC-MS**

Maize tissues were frozen in liquid $N_2$, ground to a fine powder and weighed in 60 mg aliquots. Free SA was extracted using vapor phase extraction as described previously (7). Briefly, samples were first bead homogenized in 0.3 mL 1-propanol: $H_2O$: HCl (50:25:0.4) containing 50 ng isotopically labeled [$^2H_4$]-SA (CDN Isotopes, PointeClaire, Quebec, Canada) as an internal standard (ISTD). 1 mL methylene chloride was then added and vortexed for 1 min before centrifuging at 20,000 × g for 2 min. 10 μL trimethylsilyldiazomethane was added to the supernatant and incubated at room temperature for 15 min before adding 10 μL acetic acid: methylene chloride (12:88) to neutralize the extra derivatizing reagent. The volatiles were collected with an inert filter containing 50 mg of HayeSep Q (80- to 100-mm mesh) polymer adsorbent (Sigma-Aldrich) and eluted in 200 μL methylene chloride. GC-MS analysis was conducted using an Agilent 6890 series gas chromatograph coupled to an Agilent 5973 mass selective detector (interface temperature, 250°C; mass temperature, 150°C; source temperature, 230°C; electron energy, 70 eV) as described previously (8). The gas chromatograph was operated with a DB-35MS chromatographic column (Agilent; 30 m × 0.25 mm × 0.25 μm). The sample was introduced as a splitless injection. The initial column temperature was set at 45°C for 2.25 min, then increased to 300°C with a gradient of 20°C $min^{-1}$, and held at 300°C for 5 min. The retention time for the $^{2}H_4$-SA methyl ester (ME) and endogenous SA-ME was 9.22 and 9.24 min, respectively. The diagnostic parent ion used for the MeSA quantification from endogenous SA was *m*/*z* 152, whereas MeSA [$^{2}H_4$]-SA-ME was *m*/*z* 156. Quantification of endogenous SA was calculated based upon the ratio of SA to known amounts of ISTD ([$^{2}H_4$]-SA) accounting for injection volumes, dilutions and tissue mass extracted.

**Association analysis of the *ZmGB1* locus**

We conducted candidate gene association analysis in a maize association population of 368 diverse inbred lines (9). The SNPs of ZmGB1 in this association population were downloaded from a previously released genotype dataset (9). We used KRN data from five environments including Ya'an (30°N, 103°E), Sanya (18°N, 109°E) and Kunming (25°N, 102°E) in 2009 and Wuhan (30°N, 114°E) and Kunming (25°N, 102°E) in 2010 (10), and BLUP (Best Linear Unbiased Prediction) from these five environments. The association between KRN and SNPs on ZmGB1 was established by mixed liner model corrected by population structure with $P$-value < 0.001 as a threshold (9).

**References**

1. Char SN, *et al.* (2017) An Agrobacterium-delivered CRISPR/Cas9 system for high-frequency targeted mutagenesis in maize. *Plant Biotechnol J* 15(2):257-268.
2. Brazelton VA, *et al.* (2015) A quick guide to CRISPR sgRNA design tools. *Gm Crops Food* 6(4):266-276.
3. Wu Q, Regan M, Furukawa H, & Jackson D (2018) Role of heterotrimeric Galpha proteins in maize development and enhancement of agronomic traits. *PLoS Genet* 14(4):e1007374.

4. van Wees S (2008) Phenotypic analysis of Arabidopsis mutants: trypan blue stain for fungi, oomycetes, and dead plant cells. *CSH Protoc* 2008:pdb prot4982.
5. Bommert P, Je BI, Goldshmidt A, & Jackson D (2013) The maize Galpha gene COMPACT PLANT2 functions in CLAVATA signalling to control shoot meristem size. *Nature* 502(7472):555-558.
6. Liang XX*, et al.* (2016) Arabidopsis heterotrimeric G proteins regulate immunity by directly coupling to the FLS2 receptor. *Elife* 5:e13568.
7. Schmelz EA, Engelberth J, Tumlinson JH, Block A, & Alborn HT (2004) The use of vapor phase extraction in metabolic profiling of phytohormones and other metabolites. *Plant J* 39(5):790-808.
8. Ding YZ*, et al.* (2017) Selinene Volatiles Are Essential Precursors for Maize Defense Promoting Fungal Pathogen Resistance. *Plant Physiol* 175(3):1455-1468.
9. Li H*, et al.* (2013) Genome-wide association study dissects the genetic architecture of oil biosynthesis in maize kernels. *Nat Genet* 45(1):43-50.
10. Liu L*, et al.* (2015) Genetic architecture of maize kernel row number and whole genome prediction. *Theor Appl Genet* 128(11):2243-2254.

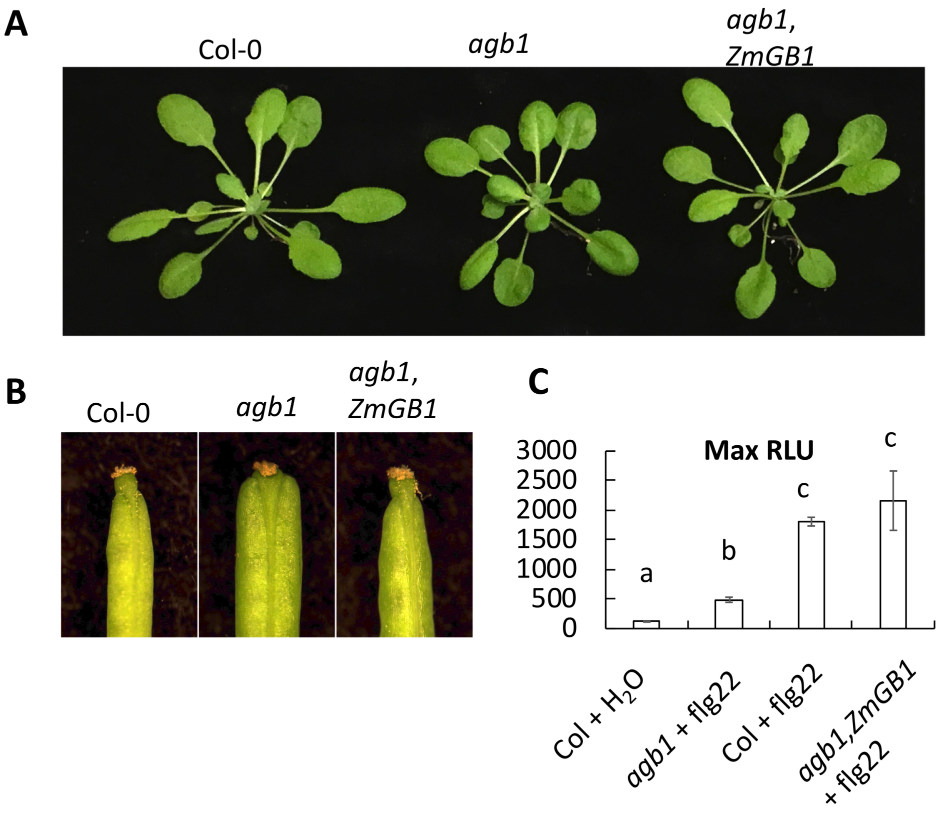


**Fig. S1. Expression of *ZmGB1* complemented both developmental and immune defects of Arabidopsis *agb1* mutants.** (A) Representative 5-weeks-old plants under short day growth conditions, note the *agb1* mutants have longer petioles and rounded leaves, whereas the *agb1* mutants complemented with *ZmGB1* resemble Col-0 wild-type. (B) Siliques of wild-type (Col), *agb1*, and *agb1* mutants expressing *ZmGB1*. (C) Leaves of the indicated genotypes were examined for flg22-induced ROS production, and peak relative luminescence unit (RLU) values were shown. Error bars indicate standard deviation; N = 6; different letters indicate significant difference with Student's t-test $p$-value $< 0.05$.

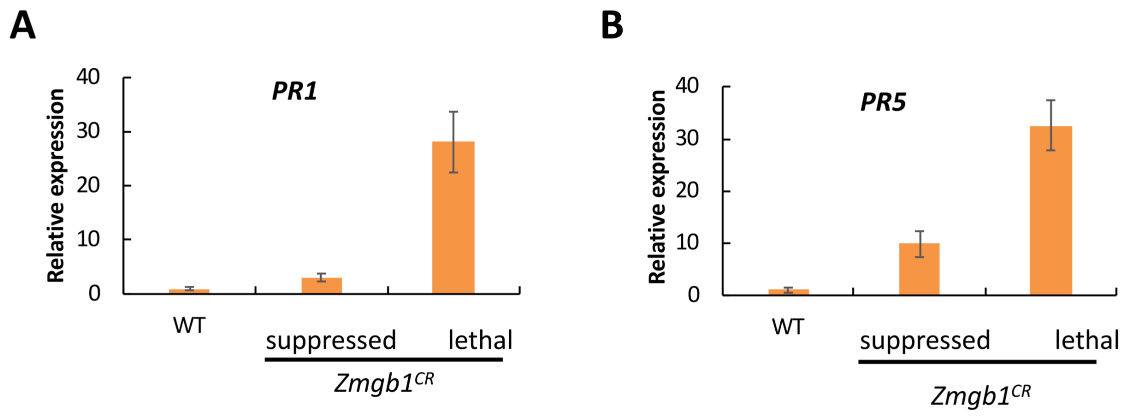


**Fig. S2**. **Induction of immune marker genes in *Zmgb1$^{CR}$* mutants was reduced in the CML103 background.** The expression of *PR1* (A) and *PR5* (B) genes was induced in the *Zmgb1$^{CR}$* lethal mutants, but their induction was lower in suppressed plants. Error bars indicate standard deviation, N =3, $p$ value < 0.001 with Student's t-test.

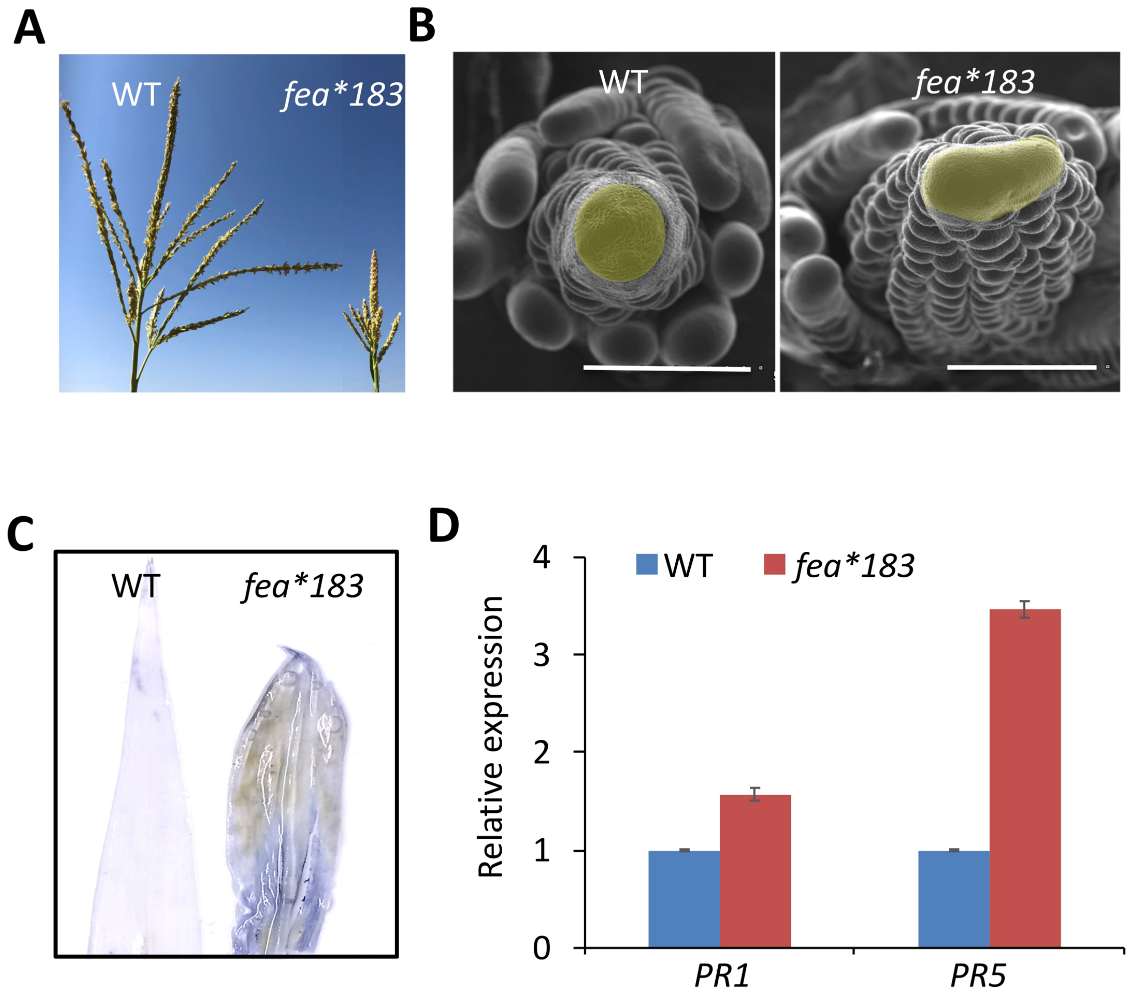


**Fig. S3. *fea*183 mutants developed inflorescence and autoimmune phenotypes.*** (A) *fea*183* mutants had compact tassels, and in (B) SEM images showed that *fea*183* mutants had enlarged tassel IMs. Scale bars = 500 μm. (C) Trypan blue staining was increased in *fea*183* leaves from 2-week-old plants. (D) *PR1* and *PR5* gene expression were up-regulated in *fea*183* compared with WT. Error bars indicate standard deviation, N =3, $p$ value < 0.001 with Student's t-test.

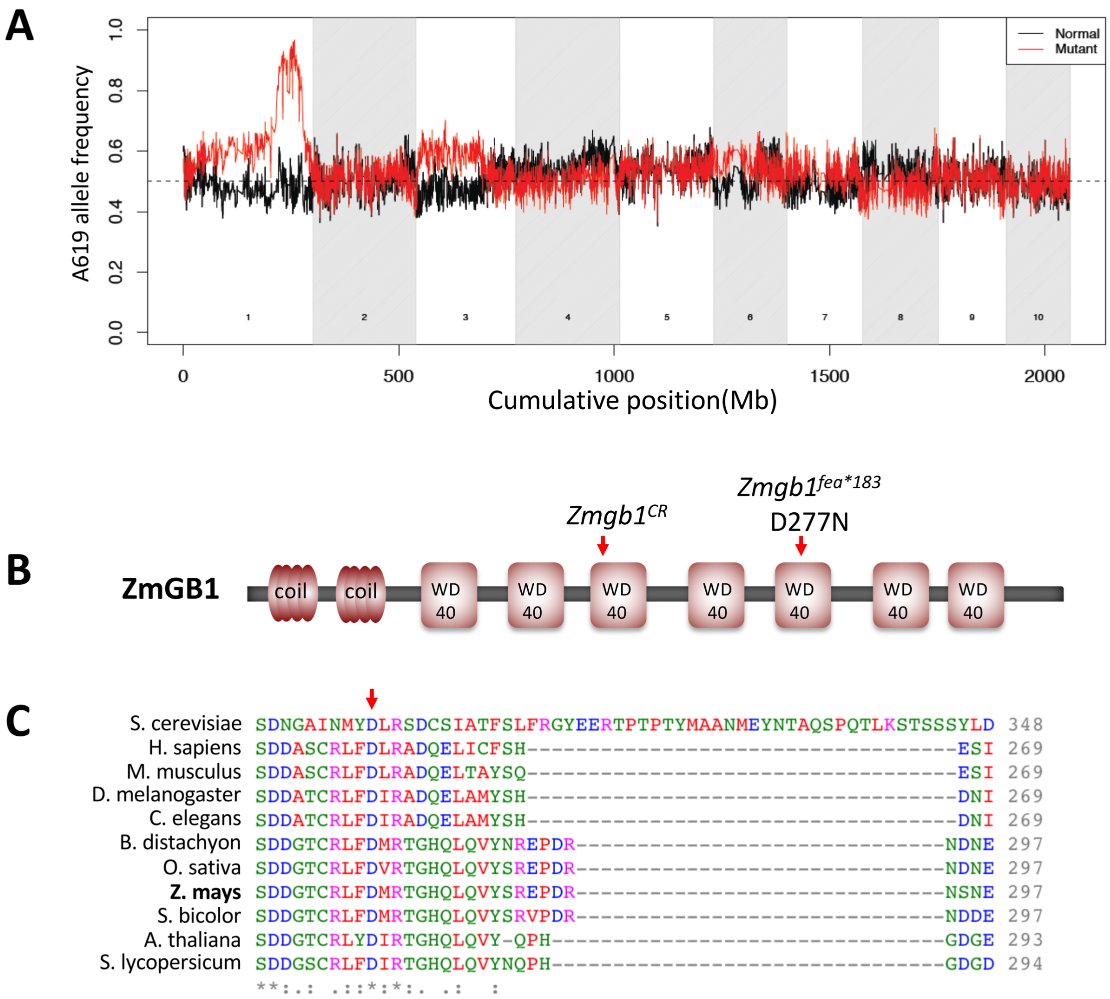


**Fig. S4. The D277N mutation in *fea*183* is in a highly conserved residue in one of the WD40 domains of ZmGB1.** (A) Bulked segregant analysis of *fea*183* using the Maize SNP50 chip revealed strong linkage at the bottom of chromosome 1. (B) Schematic of the ZmGB1 protein indicating the position of D277N in *Zmgb1*$^{*fea183}$ in one of the WD40 domains. (C) This residue is completely conserved across different species.

**Table S1** Segregation ratio of *Zmgb1*$^{CR}$*;SBP-YFP-ZmGB1* F2 family. The *pZmGB1::YFP-ZmGB1*-expressing T1 plants were backcrossed twice with *Zmgb1*$^{CR}$/+ plants in B73 background to evaluate the complementation of *Zmgb1*$^{CR}$ lethality. Chi-square was used to test the hypothesis that the transgene was able to complement the phenotype of mutants. The results showed that the observed number of lethal seedlings was not significantly different from the estimated number, suggesting that the the transgene was able to complement the phenotype of mutants. (Note, we also hypothesized that the non-transgenic control could complement the phenotype, and the chi-square test rejected this hypothesis, further confirmed our results.)

| **Event no.** | **No. of germinated seedlings** | **Surviving seedlings** | **lethal seedlings** | **chi-square p value** |
|---|---|---|---|---|
| Non-transgenic control | 32 | 24 | 8 | 0.000022 |
| 1 | 14 | 13 | 1 | 0.9296 |
| 2 | 16 | 16 | 0 | 0.3245 |
| 3 | 16 | 16 | 0 | 0.3245 |
| 4 | 34 | 31 | 3 | 0.7103 |
| 5 | 18 | 18 | 0 | 0.296 |

**Table S2** List of the primer sequences

| **Primer name** | **sequence** | **purpose** |
|---|---|---|
| ZmUBIQUITIN_F | TAAGCTGCCGATGTGCCTGCG | qRT-PCR for Ubiquitin |
| ZmUBIQUITIN_R | CTGAAAGACAGAACATAATGAGCACAG | |
| PR1_qPCR_F | TCAGCAAACAACAAACAATGG | qRT-PCR for PR1 |
| PR1-qPCR_R | GTAGTCCTGCGGCGAGTTCT | |
| PR5_qPCR_F | CGACATGAAGACCCATGCATG | qRT-PCR for PR5 |
| PR5_qPCR_R | CCTGCAAAATCCAAATCACTAGCCCA | |

| | | |
|---|---|---|
| gZmGB1-F1 | tgttGAATTCTTACTGGACACAA | Construct gRNA1 for genomic site 1, with PAM in green: GAATTCTTACTGGACACAAGGG |
| gZmGB1-R1 | aaacTTGTGTCCAGTAAGAATTC | |
| gZmGB1-F2 | gtgtGGTGGTGAATTCCCATCA | Construct gRNA2 for genomic site 2, with PAM in green: GGTGGTGAATTCCCATCAGGG |
| gZmGB1-R2 | aaacTGATGGGAATTCACCACC | |
| ZmGB1-F | TGCCTCTCAAGATGGAAGGT | PCR to genotype the target region |
| ZmGB1-R | TGTCCAACTGTAACAGCAGAC | |
| ZmGB1-MseI-F: | TGATGCTAACAGTGTGAAGT | Primer to genotype *Zmgb1*$^{fea*183}$, MseI is used to digest the PCR product |
| ZmGB1-MseI-R: | TGGTAAAAATGCAAACCTCG | |
